# Analysis of the assembly, stabilization and maturation of the multiphasic TAZ biomolecular condensates

**DOI:** 10.64898/2026.01.29.702607

**Authors:** Keren E. Shapira, Qingwei Zhu, Eden Grig, Xiaomin Chen, Yihang Jing, Xiang David Li, Haguy Wolfenson, Yoav I. Henis, Kunxin Luo

## Abstract

Phase separation is an important mechanism ensuring efficient regulation and function in Hippo signaling. Particularly, phase separation of nuclear TAZ has been demonstrated to be essential for its activity. However, the mechanisms of TAZ condensate assembly and maturation are yet undefined. Here we explored these mechanisms using FRAP with two laser beam sizes complemented by microscopy and cell biology approaches. We show that TAZ condensates are multiphasic, with a more stable core and labile periphery. TAZ initially forms small nascent clusters, likely *via* self-nucleation through the CC domain. These gradually mature into larger condensates through interaction with additional proteins *via* the WW domain. The condensates are further stabilized/activated by interaction with transcription factors and complexes including TEAD4 and P-TEFb. Of note, the ability of TAZ to form mature condensates is essential for its activities in cellular morphogenesis and tumorigenesis. Our study presents detailed mechanistic analysis of TAZ phase separation, revealing a highly dynamic nature of TAZ condensate maturation and activation.

**Teaser:** TAZ condensates grow from nascent clusters into mature condensates by interactions with transcription factors and complexes.

## INTRODUCTION

Biomolecular condensates are dynamic membraneless compartments generated by phase separation, mainly liquid-liquid phase separation (LLPS) but also liquid to gel-like or solid-like states, forming immiscible multiphase bodies which display liquid-like and gel or solid-like properties (*1–5*). They play critical functions in the regulation and control of multiple cellular processes, including cell division, signal transduction and neurodegenerative diseases (*6–8*). Numerous cellular membraneless condensates have been described, including ribonucleoprotein (RNP) particles and RNA helicase (*9, 10*), stress granules (SGs) (*11, 12*), PML (promyelocytic leukemia) bodies (*13*), and transcription-related complexes (*14–16*). As biomolecular condensates contain multiple components, they present a heterogeneous immiscible state of dense and dilute phases (*1, 3*). In particular, stress granules were reported to contain a stable core and a dynamic shell (*12*), inactivation of RNA helicase was found to induce transition of RNP particles from liquid to solid-like state (*17*), and single-component condensates of the fused in sarcoma (FUS) protein exhibit simultaneous coexistence of liquid and solid gel phases within the same condensate (*5*). Moreover, merlin, an upstream regulator of the Hippo pathway, was found to form biologically active solid-like condensates in *Drosophila* epithelia (*18*). It is therefore important to determine how specific protein-protein interactions within the biocondensates alter their physical properties and signaling activities.

We employed TAZ and Hippo signaling as a model system to investigate the dynamics of condensates formation and their functional consequences. The Hippo pathway is a highly conserved regulatory network that governs tissue growth, cell proliferation, differentiation, and organ size. It integrates multiple upstream inputs such as mechanical forces, biochemical signals and cellular stress responses to maintain homeostasis and prevent tumorigenesis. At the core of this pathway is a kinase cascade where MST1/2 phosphorylates and activates LATS1/2, leading to the phosphorylation and cytoplasmic retention or degradation of YAP/TAZ, the key transcriptional coactivators. When Hippo signaling is inactive, unphosphorylated YAP/TAZ translocate into the nucleus, binding to TEAD transcription factors to promote genes that drive cell growth, migration, and survival (*19–22*).

Recent findings highlight that phase separation plays a critical role in organizing Hippo pathway components into dynamic biomolecular condensates, which may regulate signal transduction efficiency. Upstream regulators, such as AMOT and KIBRA, undergo LLPS to form Hippo-activating condensates, which recruit MST1/2 and LATS1/2, promoting pathway activation in response to cell-cell contact and osmotic stress (*23*). In contrast, SLMAP forms Hippo-inhibiting condensates that recruit the STRIPAK phosphatase complex, leading to MST1/2 dephosphorylation and pathway inhibition (*23*). Interestingly, AMOT/KIBRA and SLMAP condensates can coalesce, counteracting STRIPAK-mediated inhibition and re-activating Hippo signaling (*23*). Additionally, core Hippo kinases, including MST1/2, SAV1, and LATS1/2, have been reported to undergo phase separation, assembling into supramolecular condensates that enhance phosphorylation efficiency and pathway activation (*24*).

Several studies have also revealed that biocondensate formation is crucial for nuclear YAP/TAZ function. Studies from our group and others have demonstrated that TAZ (*25*) or overexpression of YAP (*26*) forms nuclear condensates through LLPS. The TAZ nuclear condensates compartmentalize TEAD4 and other transcriptional elongation machinery including P-TEFb (CDK9/Cyclin T1) as well as histone modifying proteins (*25*), enhancing gene expression that drives cell proliferation and differentiation (*25–28*). Recently, the paraspeckle protein NONO has been identified as a key nuclear factor that promotes YAP/TAZ LLPS, facilitating the compartmentalization of TEAD and other cofactors required for oncogenic transcription glioblastoma (*27*). Furthermore, TAZ has been found to undergo LLPS during skeletal muscle differentiation, where it interacts with Smad7 and β-catenin to repress muscle cell differentiation (*28*). The role of LLPS in modulating YAP/TAZ nuclear activity may add a new layer of transcriptional regulation that impacts development, cancer progression, and regenerative medicine. However, the dynamic properties and assembly mechanisms of TAZ nuclear condensates have not been fully examined and their understanding is lacking. Moreover, when TAZ co-localizes in biomolecular condensates with other factors critical for its transcription activity (*e.g*., the transcription elongation factor P-TEFb), it is not clear whether P-TEFb is recruited to the TAZ condensates through direct protein-protein interactions or fusion of pre-existing TAZ and PTEF-b condensates.

In the current study, we employed biophysical and cell biological tools to explore these questions. We showed that TAZ biomolecular condensates exhibit a heterogeneous and multiphase nature and undergo a time dependent maturation process that is further stabilized by interaction with transcription factors and complexes.

## RESULTS

### TAZ condensates display multiphase properties

We have recently demonstrated that TAZ undergoes phase separation to form biocondensates both *in vitro* and *in vivo* (*25*). However, these nuclear TAZ condensates exhibit considerable heterogeneity in size and shape, and possibly in composition.

To explore the dynamic properties of these condensates and to investigate whether TAZ nuclear condensates display multiphase properties, we measured their dynamics at the center vs. the periphery of the condensate using fluorescence recovery after photobleaching (FRAP) beam-size analysis (*25, 29–31*). This method evaluates the relative contribution of lateral diffusion and exchange to the fluorescence recovery. It employs FRAP by two Gaussian beam sizes, generated by focusing the beam through ×63 (smaller beam radius) and ×40 (larger radius) objectives. If recovery is by diffusion, the characteristic fluorescence recovery time τ is the characteristic diffusion time τ_D_, which is directly proportional to the bleached area (τ_D_ = ω^2^/4*D*, where ω is the beam Gaussian radius and *D* is the lateral diffusion coefficient). Therefore, a τ(×40)/τ(×63) ratio equal to the ratio between the bleached areas (2.28 in our FRAP setup; see (*30*)), indicates that the recovery occurs by diffusion. For phase separation, the expectation is for recovery by lateral diffusion within each phase (within and outside the condensate), as each phase behaves like a liquid. On the other hand, when the recovery is by exchange between a relatively immobile fluorescent structure and the surroundings, τ reflects the chemical relaxation time, which does not depend on the beam size [i.e., τ(×40)/τ(×63) = 1]. Such a behavior is expected for a relatively immobile condensate in a gel/solid phase, as the contribution of diffusion within the condensate to the fluorescence recovery would be negligible.

In accord with our earlier study (*25*), GFP-TAZ expressed in HeLa cells formed visible condensates (nuclear puncta) at 24 h post-transfection (Fig. 1A). A typical FRAP curve obtained at the center of a condensate is shown in Fig. 1B, and the averages of multiple FRAP experiments using ×63 and ×40 objectives are depicted in Fig. 1C-E. These studies demonstrate that GFP-TAZ within condensates recovers by lateral diffusion, as indicated by a τ(×40)/τ(×63) ratio not significantly different from the beam-size ratio, 2.28 (Fig. 1E, F). The GFP-TAZ diffusion coefficient (*D*) in nuclear puncta calculated from these values based on the area bleached by the Gaussian beam is 0.09 μm^2^/s, very close to the value measured by us earlier and to *D* measured for other proteins in nuclear condensates (*10, 11*). Analogous FRAP studies in regions with diffuse GFP-TAZ [either in the cytoplasm (Cyto) or the nucleoplasm (Nuc)] also yielded τ(×40)/τ(×63) ratios typical of recovery by diffusion, albeit with much faster diffusion rates (*D* = 2.2 and 1.8 μm^2^/s for diffusion in the cytoplasm or nucleoplasm, respectively; Fig. 1D, F). Thus, fluorescence recovery occurs by diffusion both at regions with diffuse GFP-TAZ distribution (*e.g*., nucleoplasm) and in condensates, in line with phase separation. Of note, the recovery was nearly complete in the diffuse regions as indicated by the mobile fraction (*R_f_*) approaching 1, but was much lower in nuclear condensates (Fig. 1C). These results confirm that TAZ forms nuclear phase separated condensates as we have reported earlier (*25*).

**Fig. 1.**
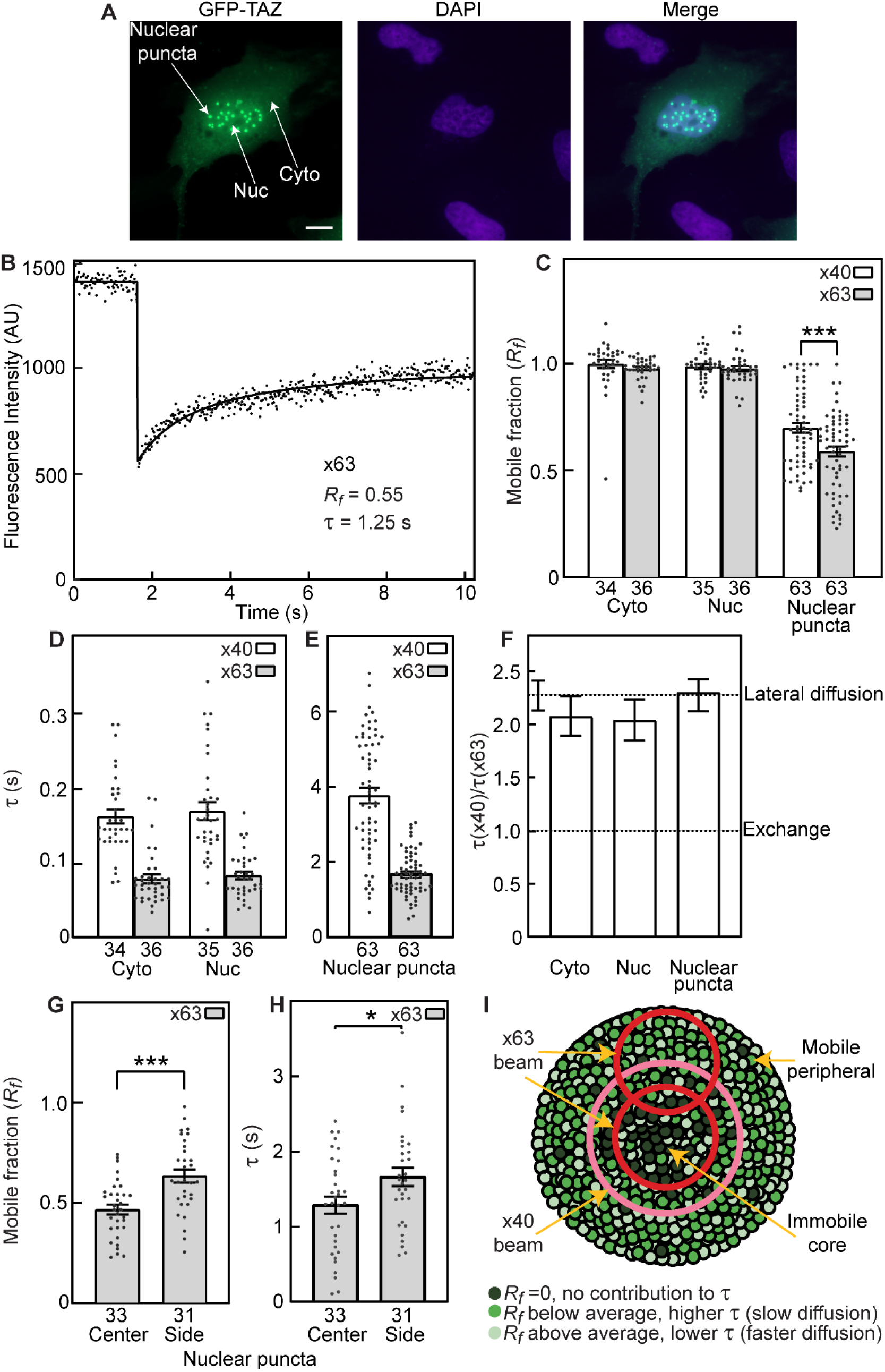
FRAP beam-size analysis reveals multiphase properties of GFP-TAZ-WT condensates. HeLa cells were transfected with GFP-TAZ-WT and subjected to FRAP beam-size analysis studies 24 h later. **(A)** Fluorescence images of GFP-TAZ condensates taken with the FRAP microscope. Arrows indicate representative areas of measurement: visible nuclear condensates (nuclear puncta), and diffuse cytoplasmic (Cyto) or nucleoplasmic (Nuc) distributions. Similar results were obtained from 3 independent transfections. **Bar**, 10 μm. **(B)** A typical FRAP curve (×63 objective) of GFP-TAZ-WT in a nuclear condensate. Solid line, best fit (nonlinear regression) to the lateral diffusion equation**. (C-F)** FRAP beam-size analysis in nuclear condensates and in diffuse cytoplasmic (Cyto) or nucleoplasmic (Nuc) regions. Bars are mean ± SEM of multiple independent measurements, each on a different cell (the number of measurements is depicted underneath each bar in panels C-E, G, H). The studies employed ×40 or ×63 objectives, focusing the beam to larger (×40) or smaller (×63) areas, with a beam size ratio of ω^2^(×40)/ω^2^(×63) = 2.28 ± 0.15 (n = 59 independent measurements). SEM for the *τ* ratios (F) were calculated from the *τ* values in panels D and E, using bootstrap analysis (1,000 bootstrap resampling values). In (C), the pairs of the *R_f_* values obtained with the ×40 or ×63 objectives for each specific condition were compared. Asterisks indicate significant differences between the pairs indicated by brackets (***, *P*<10^−3^; Student’s two-tailed *t*-test). In (F), all τ(×40)/τ(×63) ratios were not significantly different from the 2.28 beam-size ratio (Student’s two-tailed *t*-test, comparing each τ ratio to the beam-size ratio), suggesting FRAP by lateral diffusion in all cases. **(G, H)** The center *vs*. periphery of GFP-TAZ-WT nuclear condensates display different *R_f_* (G) and τ (H) values. The studies were performed with the smaller beam size (×63 objective), focusing the beam at the condensate center or periphery (designated “side”). Asterisks indicate significant differences between the same parameter at the center and at the periphery (*, *P*<0.05; ***, *P*<10^−3^; Student’s two-tailed *t*-test). **(I)** Scheme depicting the principle of FRAP with different beam sizes on condensates. The principle is shown for TAZ but holds also for other molecules. TAZ molecules in condensates display heterogeneous interactions with multiple components (other TAZ molecules, other proteins, RNA). For simplicity, three classes of TAZ molecules experiencing different degrees of mobility-restricting interactions are depicted (dark, medium and light green for high, medium and low level of mobility restriction). The condensate center is enriched with molecules subject to a high level of mobility-restricting interactions, while the periphery has a high percentage of molecules with medium and low mobility restriction. Thus, FRAP with a small beam size (×63) at the center yields low *R_f_* and faster τ, since τ at the center is contributed largely by the low-restricted population, while FRAP at the periphery shows a higher *R_f_* and slower τ. Moreover, FRAP with the larger beam (×40) at the center covers a larger area, which contains the immobile core plus a large population of molecules with medium mobility restriction, resulting in a higher *R_f_* and a slower τ.

Interestingly, we noticed two phenomena which indicate that the GFP-TAZ nuclear condensates have a multiphase nature, in line with the suggested heterogenous interactions in biocondensates (*1–3*). First, the mobile fraction at the center of the condensate (Fig. 1C, nuclear puncta) was significantly lower for FRAP with the smaller beam size (×63 objective) than with the larger beam size (×40 objective; *R_f_* = 0.58 *vs*. 0.70, respectively). This indicates that most of the immobile TAZ molecules reside at the center of the condensate. Thus, when the smaller ×63 beam is employed, it covers mainly this immobile core plus a small fraction of the mobile periphery, while the ×40 objective, which bleaches a larger area around the center, encompasses a larger fraction of the mobile periphery, leading to a higher *R_f_* (see scheme of the experimental design in Fig. 1I). Second, FRAP with the smaller beam size (×63 objective) at different regions of the GFP-TAZ nuclear puncta (center *vs*. periphery; Fig. 1I) yielded different *R_f_* values, with *R_f_* at the center being significantly lower than at the periphery (Fig. 1G). Moreover, this lower *R_f_* in the center is accompanied by a shorter τ (Fig. 1H; see scheme in Fig. 1I). Of note, the τ values measured are weighted averages of all mobile GFP-TAZ complexes in the bleached region, with no contribution from the immobile core at the center of the condensate. Thus, focusing the ×63 beam at the periphery includes a higher contribution to τ by TAZ molecules displaying both medium and low levels of mobility-restricting interactions, while FRAP at the center reflects a higher fraction of the immobile population, which does not contribute to the τ measurement. Concomitantly, for FRAP at the center, some of the slow-diffusing GFP-TAZ molecules may be effectively immobilized by interactions with the immobile core, leaving only the faster-diffusing molecules to contribute, yielding a faster τ value. Taken together, TAZ nuclear condensates display a heterogeneous and multiphase nature, as reported for several other biocondensates (*1–3, 32, 33*), and their assembly likely involves dynamic processes.

### TAZ biomolecular condensates display time-dependent maturation

We next conducted FRAP beam-size analysis on GFP-TAZ mutants displaying enhanced or defective visible puncta formation (Fig. 2A). First, we studied TAZ-S89A, a constitutively active TAZ mutant resistant to LATS1/2 inhibition (*25, 34*), which we have shown to form large nuclear condensates (Figs. 2A, 3A and (*25*)). The results of the FRAP experiments (Fig. 2B-E) demonstrated that *R_f_* of GFP-TAZ-S89A in nuclear condensates, measured with the larger beam (×40 objective), is lower than that of TAZ-WT, becoming as low as *R_f_* measured with the ×63 objective (Fig. 2B). This may suggest that the TAZ-S89A condensates contain a larger immobile core, such that when the larger ×40 beam is focused at the TAZ-S89A condensate center, it predominantly covers this immobile core, in a similar manner to how the ×63 objective covers the immobile core of TAZ-WT and TAZ-S89A condensates (see scheme in Fig. 1I). This interpretation is supported by FRAP studies with the smaller (×63) beam size focused at the center vs. periphery of the condensates (Fig. 2D, E), which show reduced mobility (lower *R_f_*) and faster τ at the center relative to the periphery, in line with the results obtained for TAZ-WT (see Fig. 1G, H). These results provide an additional support to the notion that TAZ condensates contain a stable center core and a more labile periphery, consistent with time-dependent maturation that initiates from the center of the condensates and progresses outwards.

**Fig. 2.**
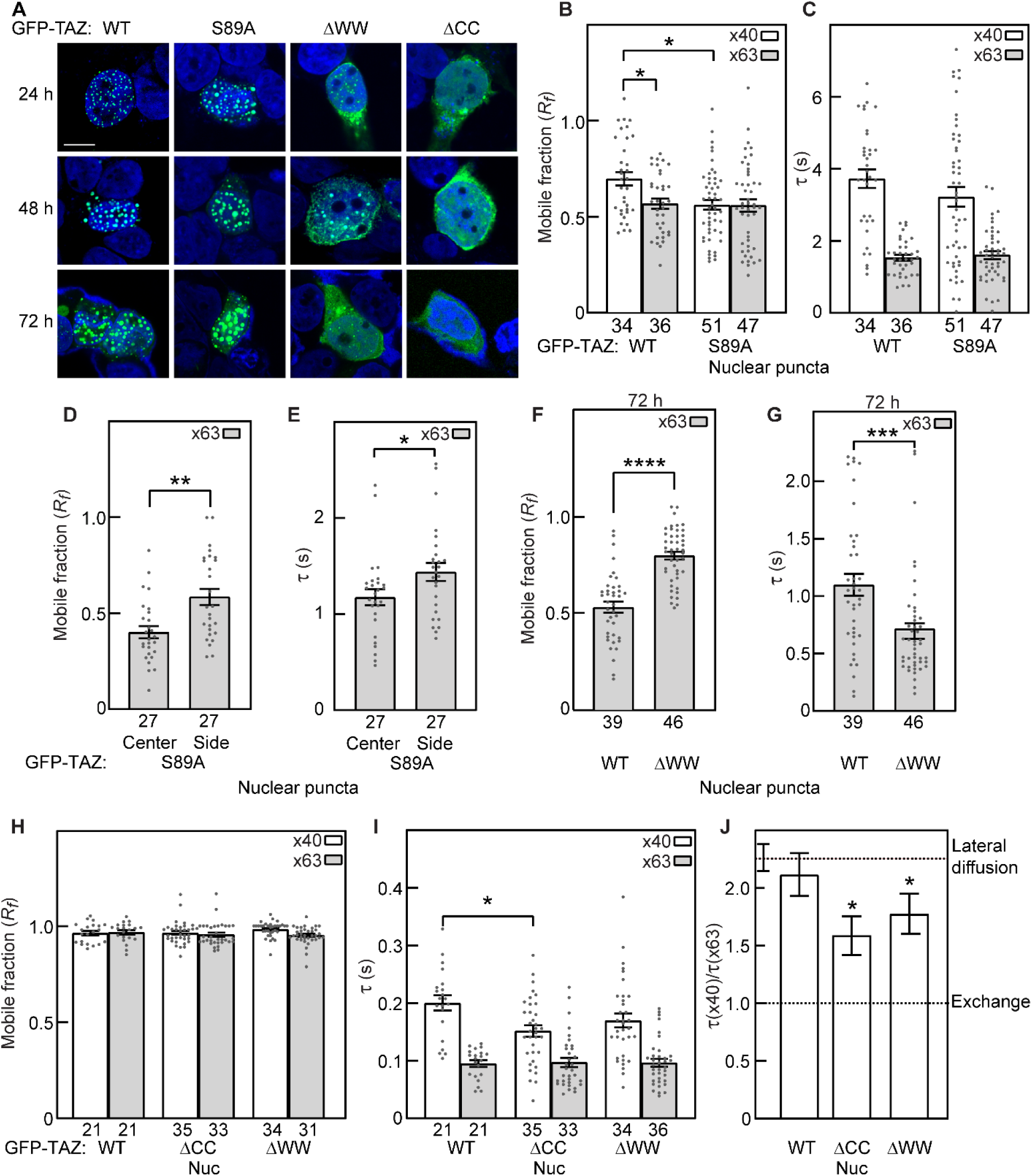
FRAP studies of GFP-TAZ mutants. HeLa cells were transfected with WT or mutant GFP-TAZ and analyzed as in Fig. 1. **(A)** Representative confocal images (Zeiss LSM 710 confocal microscope). Green, GFP-TAZ variants; blue, DAPI. **Bar**, 10 μm. **(B-J)** FRAP experiments. Bars are mean ± SEM of multiple independent measurements, whose numbers are shown beneath the bars. **(B, C)** FRAP studies (24 h post-transfection) in nuclear condensates of GFP-TAZ-S89A and GFP-TAZ-WT. Asterisks indicate significant difference between the pairs indicated by brackets (*, *P*<0.05; one-way ANOVA and Tukey’s post hoc test). **(D, E)** GFP-TAZ-S89A displays lower *R_f_* and shorter τ values at the center of nuclear condensates. FRAP studies employed the smaller beam size (×63 objective) to measure *R_f_* (panel D) and τ (panel E) at the center *vs*. side of GFP-TAZ-S89A nuclear condensates. Asterisks indicate significant differences between the pairs indicated by brackets (*, *P*<0.05; **, *P*<0.01; Student’s two-tailed *t*-test). **(F, G)** GFP-TAZ-ΔWW condensates are more labile than GFP-TAZ-WT condensates. GFP-TAZ-ΔWW required longer (72 h) than TAZ-WT to form visible condensates. FRAP measurements employed the ×63 objective to fit within these condensates; GFP-TAZ-WT condensates at 72 h post-transfection were measured as control. Asterisks indicate significant differences between the pairs indicated by brackets (***, *P*<10^−3^; ****, *P*<10^−4^; Student’s two-tailed *t*-test). **(H-J)** FRAP beam-size analysis of GFP-TAZ-ΔWW and -ΔCC in nucleoplasmic regions (Nuc) lacking visible condensates. Experiments were conducted 24 h post-transfection with the ×40 and ×63 objectives, as in Fig. 1C-F. No significant differences were obtained between the *R_f_* values of the ×40 or ×63 objectives (H), which were all close to 1. The τ(×40) or τ(×63) values of the TAZ mutants were also close to those of TAZ-WT (I), the only difference being a somewhat lower τ(×40) for TAZ-ΔCC (*, *P*<0.05; one-way ANOVA and Tukey’s post hoc test). **(J)** Bootstrap analysis of the τ(×40)/τ(×63) ratio relative to the beam size ratio (2.28). SEM values for the *τ* ratios were calculated from the *τ* values shown in panel I, using 1,000 bootstrap resampling values. While the τ ratio of GFP-TAZ-WT was similar to the beam-size ratio, the τ ratios of the ΔCC and ΔWW mutants deviated significantly from beam size ratio (*, *P*<0.05; Student’s two-tailed *t*-test).

**Fig. 3.**
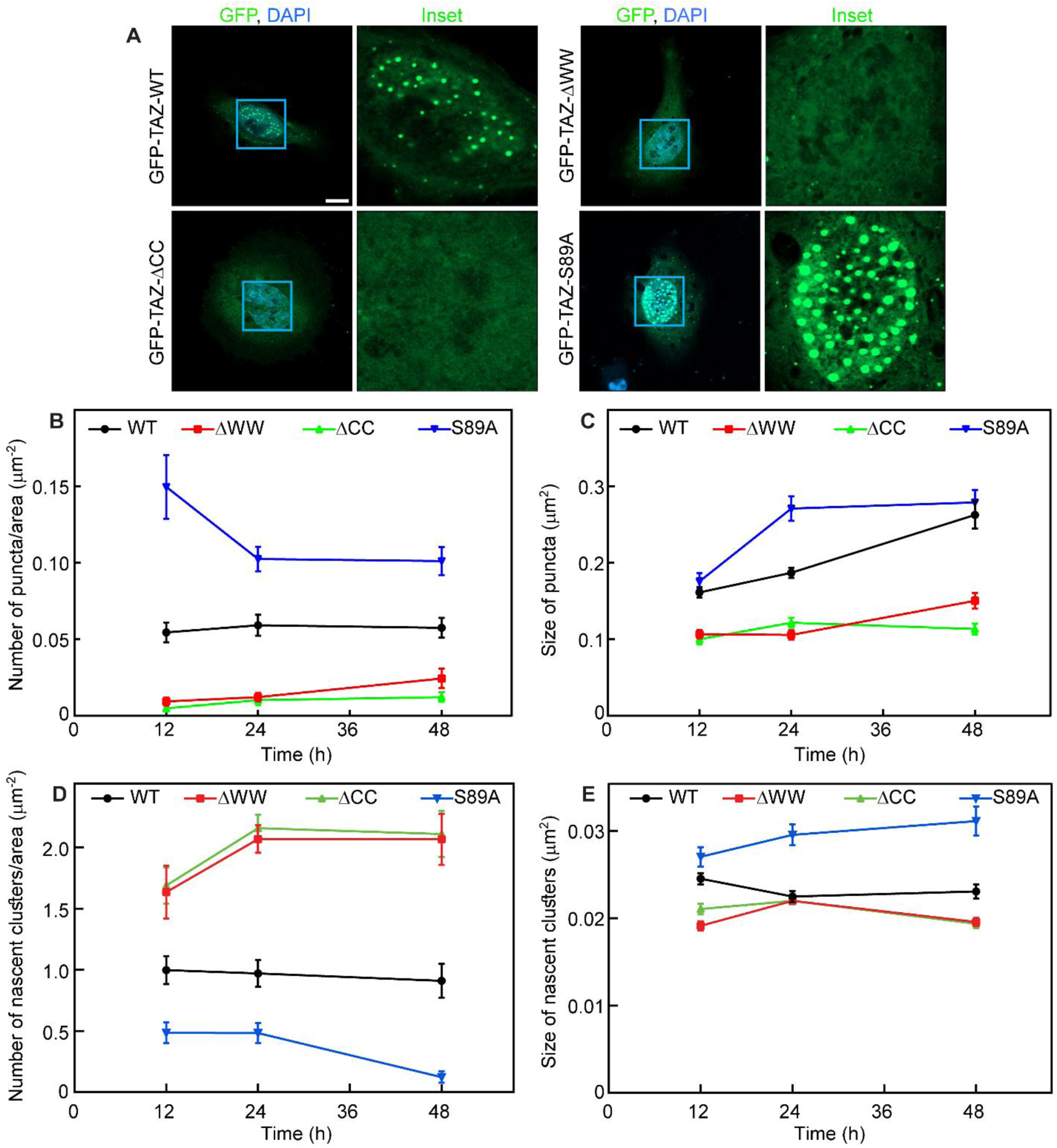
Time course analysis of visible puncta and of nascent clusters formed by GFP-TAZ and mutants. HeLa cells were transfected with either WT or mutant GFP-TAZ as indicated. At the indicated times, they were fixed, mounted and subjected to super-resolution microscopy analysis. Puncta or clusters in the nuclei of multiple cells were counted by Fiji (Image J), normalizing per nuclear area. **(A)** Representative super-resolution images of GFP-TAZ variants at 24 h, obtained using Zeiss LSM800 Airyscan microscope. For each mutant, the green and blue channels (GFP and DAPI) are depicted in the left frame, and an enlarged inset of the green channel (x3.61 magnification) is shown on the right. Similar results were obtained in 5 independent transfections. **Bar**, 10 μm. **(B, C)** Mean ± SEM of the density (number per area) of visible puncta (B) and average puncta size (C) at different time points post transfection. To capture only visible condensates, a lower threshold of 0.06 μm^2^ was set). The original data points and statistics of panels B, C are depicted in fig. S1. **(D, E)** Mean ± SEM of the density of nascent clusters (D) and their size (E) at the same time points. Nascent clusters were defined as occupying areas between 0.008 μm^2^ and 0.05 μm^2^. The original data points and statistics are shown in fig. S2.

Next, we conducted analogous studies on two GFP-TAZ mutants defective in puncta formation: GFP-TAZ-ΔWW, which lacks the WW domain and has a considerably reduced ability to form condensates in cells, and GFP-TAZ-ΔCC, where the CC domain is deleted and visible puncta formation is abolished (*25*). Consistent with our previous report, the ΔCC mutant failed to form visible puncta in cells even at 72 h post-transfection (Fig. 2A), and was therefore used as a negative control. The ΔWW mutant formed much smaller puncta at 48 h and required 72 h to form visible puncta, which were still smaller than those of TAZ-WT (Figs. 2A and 3A, C). FRAP measurements on the visible nuclear puncta formed by GFP-TAZ-ΔWW at 72 h (Fig. 2F, G) were conducted using the smaller beam size (×63 objective), which can fit inside the smaller condensate. The *R_f_* of GFP-TAZ-ΔWW in condensates was significantly higher than that of GFP-TAZ-WT at 72 h (Fig. 2F), indicating that a lower fraction of GFP-TAZ-ΔWW resides in the immobile core relative to GFP-TAZ-WT. This was accompanied by faster recovery (shorter τ) of TAZ-ΔWW as compared to TAZ-WT (Fig. 2G). These results suggest that the condensates formed by GFP-TAZ-ΔWW are significantly smaller, more labile and less stable than those of GFP-TAZ-WT.

### The nascent nucleoplasmic clusters of TAZ-ΔWW and TAZ-ΔCC are more labile than those of TAZ-WT

Small (nascent) clusters, referred to as higher order oligomers or pre-condensate clusters, were reported to form when the phase-separating proteins were below the threshold concentration for condensate formation (*35, 36*). To evaluate whether the TAZ condensates initiate as nascent clusters, we conducted FRAP beam-size studies on TAZ-WT and its mutants in the nucleoplasm (diffuse-labeled areas) at 24 h post-transfection (Fig. 2H-J). The *R_f_* values of the ΔWW and ΔCC mutants were similar to those of GFP-TAZ-WT (Fig. 2H). However, while the τ(×40)/τ(×63) ratio of GFP-TAZ-WT in the nucleoplasm fitted FRAP by lateral diffusion (2.28), those of the ΔWW and ΔCC mutants showed significant downward deviations (1.8 and 1.6, respectively; Fig. 2J), due to shorter τ(×40) values (Fig. 2I). This indicates that unlike TAZ-WT, FRAP of the ΔCC and ΔWW mutants in the nucleoplasm occurs not only by diffusion, but also contain a contribution of exchange, in line with higher lability. A possible explanation is that at the 24 h time point, the concentration of the ΔWW or ΔCC mutants in the nucleoplasm are insufficient for visible puncta formation, but they can form nascent GFP-TAZ nucleation centers (pre-condensate clusters), which are below the resolution of light microscopy. Indeed, this notion is supported by super-resolution microscopy studies (Zeiss Airyscan, providing ∼120 nm resolution) on cells expressing the TAZ mutants (Fig. 3). If these nascent puncta were stable, their exchange rate with the diffuse nucleoplasmic population would be significantly slower than the lateral diffusion rate, and only the contribution of the faster process (diffusion) to recovery would be observed, as in the case of GFP-TAZ-WT. However, if the exchange rate becomes faster and approaches that of the lateral diffusion rate for labile nascent puncta, as in the case of the ΔWW and ΔCC TAZ mutants, the rates of exchange and diffusion would be close, resulting in a significant contribution of both exchange and diffusion to the measurement.

To test this hypothesis, we conducted super-resolution microscopy studies on HeLa cells expressing GFP-TAZ (WT or mutant) as a function of time, and quantified the number and size of both visible and nascent puncta (Fig. 3; for statistics and data spread, see Supplementary Figs. 1, 2) using Fiji (Image J). Nascent nucleation clusters visible only at super-resolution were defined by setting a low threshold at 0.1 μm diameter (circle area of 0.008 μm^2^) and an upper threshold at 0.24 μm diameter (0.05 μm^2^ area). Puncta above the latter threshold were defined as visible condensates. As shown in Fig. 3A-C and fig. S1, GFP-TAZ-WT and GFP-TAZ-S89A formed visible puncta (condensates) at all time points tested, with a higher density and a larger size (especially at 24 h) for the constitutively active TAZ-S89A, while TAZ-ΔWW formed very few and much smaller visible puncta, and TAZ-ΔCC barely formed any. The density (number per nucleus area) for all TAZ variants was rather constant. On the other hand, the size of the visible puncta formed by TAZ-WT and TAZ-S89A increased with time (Fig. 3C), in line with time-dependent maturation of the condensates.

In contrast, analysis of the super-resolution images for nascent cluster formation demonstrated a highly different pattern (Fig. 3D, E). At all-time points, the number of nascent nuclear clusters per area was highest for the TAZ mutants defective in condensate formation (ΔWW and ΔCC), significantly lower for TAZ-WT, and lowest for TAZ-S89A (Fig. 3C and fig. S2A-C). This pattern is a mirror image of the density of the larger visible condensates as a function of time (Fig. 3B), and is also reflected in the average size of the initial clusters, which was smaller for TAZ-ΔWW and TAZ-ΔCC (especially at 12 and 48 h), and highest for TAZ-S89A (Fig. 3D and fig. S2D-F). These results suggest that formation of TAZ biomolecular condensates initiates as small nascent clusters, which are subsequently stabilized and enlarged from the center to the periphery through recruiting interacting proteins. This maturation process appears to depend on the CC and WW domains, as both the ΔCC and ΔWW mutants can form nascent clusters to a high degree (Fig. 3D, E), but either fail to grow further into visible condensates even at 72 h (TAZ-ΔCC), or form very few condensates at 48 h which mature to smaller condensates at 72 h (TAZ-ΔWW) (Fig. 3A-C and fig. S1). Thus, the CC and WW domains of TAZ likely mediate protein-protein interactions at the early stages of condensate formation from nascent clusters. While the CC domain is essential for condensate formation, the WW domain plays a facilitative role for condensate stabilization and maturation.

To further explore how the TAZ CC domain might facilitate self-nucleation, we employed AlphaFold2 (*37*) to predict the structure of the TAZ CC domain and whether it can form higher-order aggregates. As shown in Fig. 4A, the CC domain not only exists as a dimer as shown earlier by biochemical methods (*25*), but more interestingly, is predicted to form a tetrameric assembly. This tetrameric configuration requires Cys262 (C262), which is positioned to form intermolecular disulfide bonds that could facilitate higher-order aggregation. To test this hypothesis, we expressed and purified the TAZ CC domain fused to a Thioredoxin (TrxA) tag (TAZ-CC) and characterized its oligomerization state using size-exclusion chromatography coupled with multi-angle light scattering (SEC-MALS) in the presence or absence of the reducing agent dithiothreitol (DTT). The results (Fig. 4B) demonstrated that treatment with 2 mM DTT significantly impaired oligomerization of the CC domain. More importantly, when C262 was mutated to Ser (C262S), the mutated TAZ-CC domain construct behaved predominantly as a monomer. Taken together, these data suggest that C262-mediated disulfide bonding promotes the self-oligomerization of the TAZ-CC domain, contributing to its initial self-nucleation activity.

**Fig. 4.**
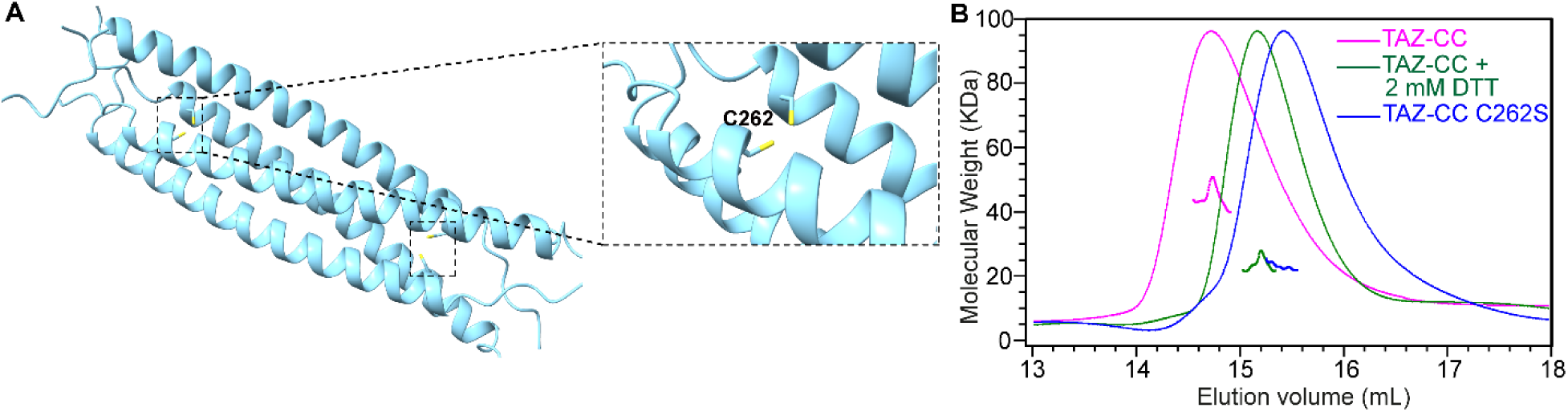
Dimers of the TAZ CC domain form higher order aggregates *via* C262-mediated disulfide bond. (**A**) Structure of a tetrameric assembly formed by the TAZ CC domain as predicted by AlphaFold2. Two sets of TAZ-CC dimers form a tetrameric structure *via* an intermolecular disulfide bond established by C262. (**B**) FPLC-coupled static light scattering showing the oligomerization states of TAZ-CC C262S mutant (blue) and WT TAZ-CC in the absence (magenta) or presence (green) of 2 mM DTT (50 μM loading concentration). The fitted molecular weights are shown in the graph.

### TEAD is recruited into and stabilizes TAZ condensates

TAZ binds to the DNA-binding cofactor TEAD and through this interaction, gets recruited to specific promoter DNA sequences. Our previous work has shown that TEAD can be recruited into TAZ nuclear condensates *via* its physical interaction with TAZ (*25, 38*). As shown in Figure 4A, singly expressed TEAD4 was homogeneously distributed in the nucleus. Upon coexpression with TAZ-WT, TEAD4 became highly enriched within distinct TAZ nuclear puncta. Deletion of the TEAD-binding domain (TAZ-ΔTB) in TAZ abrogated TEAD4 recruitment into the TAZ puncta, confirming the requirement for TAZ-TEAD interaction. Interestingly, GFP-TAZ-ΔTB formed large visible condensates at 24 h as visualized under confocal microscopy (Fig. 5A) and super resolution microscopy (Fig. 5B). These GFP-TAZ-ΔTB condensates exhibited dynamic properties different from those of GFP-TAZ-WT in both the *R_f_* and τ values (Fig. 5C-E). Unlike GFP-TAZ-WT, the *R_f_* values of GFP-TAZ-ΔTB measured with the two beam sizes (×40 and ×63 objectives) were similar (Fig. 5C). This observation correlates with the larger condensates formed by the ΔTB mutant at 24 h (Fig. 5G), indicating that the immobile core in GFP-TAZ-ΔTB condensates is larger, and therefore occupies most of the area illuminated by the ×40 objective when focused at the center of the GFP-TAZ-ΔTB condensate. However, the large condensates formed by GFP-TAZ-ΔTB at 24 h are not stable, as evidenced by the decrease in puncta size from 24 h to 48 h (Fig. 5G), as well as by the marked reduction in the number per unit area of TAZ-ΔTB condensates between 12 h and 48 h (Fig. 5F). This differs from the stable number of GFP-TAZ-WT condensates and their steady growth in size with time (Fig. 3B, C). The instability of GFP-TAZ-ΔTB relative to GFP-TAZ-WT condensates is supported by the faster recovery times [τ(×40) and τ(×63)] of the ΔTB condensates (Fig. 5D), suggesting that GFP-TAZ-ΔTB molecules miss some restrictions on their diffusion, most likely due to the loss of interactions conferred by the TEAD binding domain. Thus, binding of TEAD may stabilize TAZ condensates.

**Fig. 5.**
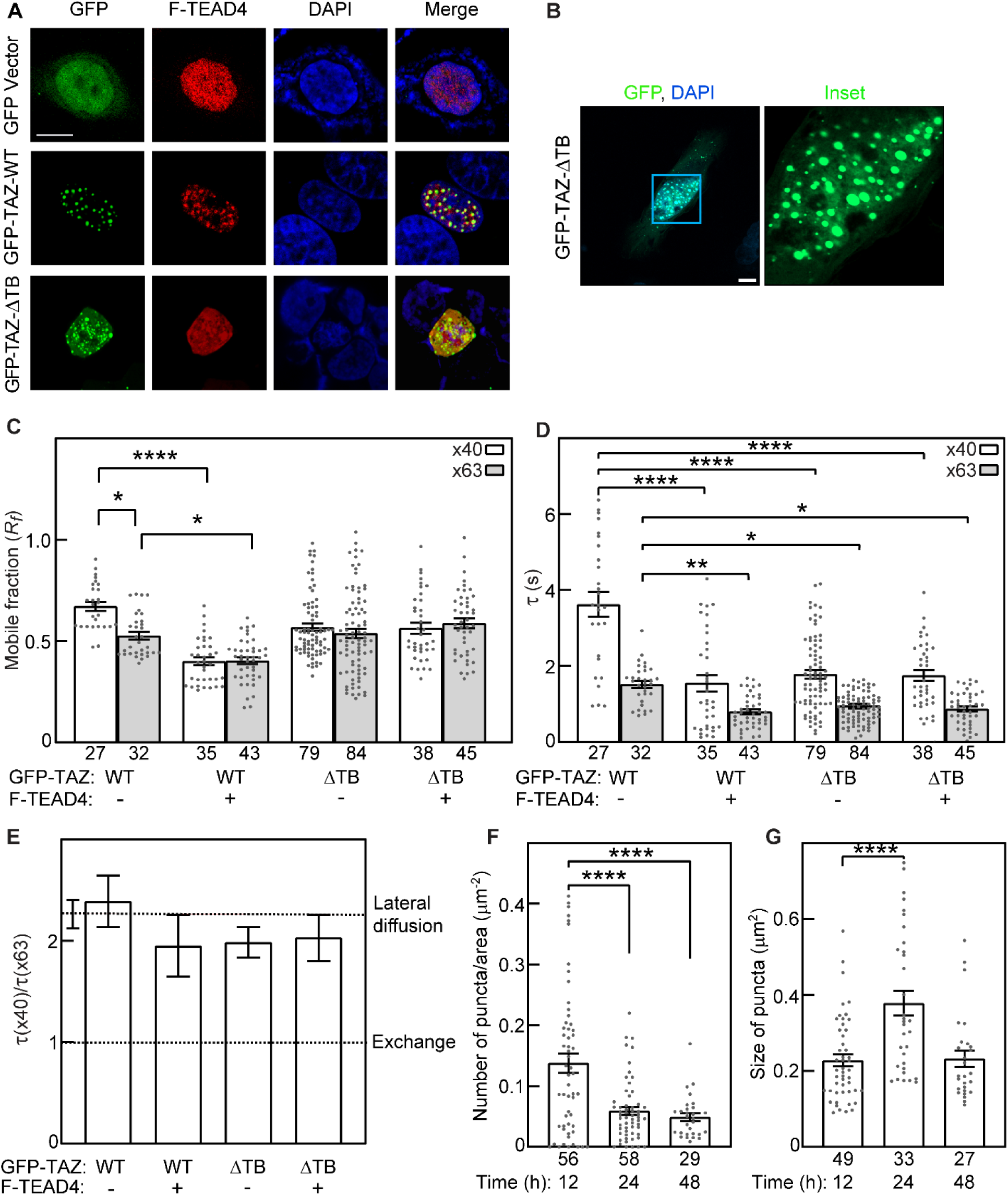
TEAD binding contributes to the stability of GFP-TAZ condensates. HeLa cells were transfected by GFP-TAZ-WT or -ΔTB, without or with Flag-TEAD4 (F-TEAD4). Bars are mean ± SEM of multiple independent measurements (whose numbers are shown under the bars), each on a different cell. **(A)** Representative images (Zeiss LSM 710 confocal microscope). **Bar**, 10 μm. **(B)** Representative super-resolution images (Zeiss LSM800 Airyscan microscope, employed for the studies in F, G). The frame on the left shows the green and blue channels; the right frame depicts an enlarged inset of the green channel (x3.61 magnification). Similar results were obtained in 5 independent transfections. **Bar**, 10 μm. **(C-E)** Average *R_f_* (C), τ (D) and τ ratio (E) of FRAP beam-size analysis on condensates of GFP-TAZ variants 24 h post-transfection as in Fig. 1. In (C, D), asterisks indicate significant differences between the pairs indicated by brackets (*, *P*<0.05; **, *P*<0.01; ****, *P*<10^−4^ ; one-way ANOVA and Tukey’s post hoc test). **(E)** Bootstrap analysis of the τ(×40)/τ(×63) ratio relative to the beam size ratio (2.28). The SEM values for the *τ* ratios were calculated from the *τ* values shown in panel C, using 1,000 bootstrap resampling values. This analysis showed no significant differences from the ratio expected for recovery by lateral diffusion. **(F-G)** Quantification of the number (F) and size (G) of GFP-TAZ-ΔTB visible nuclear puncta. Cells were transfected with GFP-TAZ-ΔTB, and incubated for 12, 24 or 48 h. Super-resolution microscopy analysis was performed as in Fig. 3. The number and size of visible puncta were analyzed by Fiji (Image J), with a lower threshold set to 0.06 μm^2^. Puncta numbers were normalized per nuclear area. Significant differences between pairs were evaluated by one-way ANOVA and Tukey’s post hoc test (****, *P*<10^−4^).

To further analyze the outcome of TEAD binding, we investigated the effects of TEAD4 overexpression on the properties of GFP-TAZ-WT condensates at 24 h by FRAP beam-size analysis (Fig. 5C-E). Coexpression of Flag-TEAD4 with GFP-TAZ-WT resulted in condensates with *R_f_* values significantly lower than those of TAZ-WT, and reduced *R_f_* with the ×40 objective to the level obtained with the smaller beam size (Fig. 5C). Both effects are consistent with an enlarged immobile core and increased stabilization. Of note, the *R_f_* values of GFP-TAZ-ΔTB were unaffected by expression of TEAD4 (Fig. 5C), reinforcing the notion that the stabilization of TAZ-WT condensates requires TEAD association. The stabilization of GFP-TAZ-WT condensates by TEAD is further supported by the shift of TAZ-WT recovery [τ(×40) and τ(×63)] to shorter times (Fig. 5D), a phenomenon likely to reflect a loss of the contribution of slower-diffusing molecules to τ due to their recruitment to the immobile core. As in the case of the *R_f_* values, the τ values of the GFP-TAZ-ΔTB mutant were unaffected by TEAD4, demonstrating the specificity of the effect. We conclude that although TAZ condensates can form without TEAD, TEAD binding is important for their stabilization.

### P-TEFb components coalesce with TAZ condensates and alter their dynamics and physical properties

Our previous study has shown that the P-TEFb transcription elongation factor, consisting of CDK9 and Cyclin T1 (CycT1), is present in TAZ nuclear condensates and likely plays an important role in TAZ transcription activity (*25*). Interestingly, CycT1 itself is known to undergo phase separation and initiate the formation of P-TEFb condensates critical for transcription elongation (*16*). To determine how CycT1 localizes into TAZ condensates, HeLa cells expressing mCherry-CycT1 alone or together with GFP-TAZ-WT were examined by fluorescence confocal microscopy and live-cell confocal imaging, where cells were imaged over 300 cycles within a 5-min period to capture rapid nuclear dynamics. As shown in Fig. 6A and movie S1, singly expressed mCherry-CycT1 localized primarily to the nucleus and exhibited a diffuse distribution with occasional faint enrichment in certain nuclear regions, suggestive of weak or nascent condensate formation. However, no distinct puncta or clear compartmentalization were detected. Coexpression with GFP-TAZ-WT progressively recruited CycT1 into more compact and distinct nuclear puncta that co-localized with GFP-TAZ. Merged fluorescence signals showed strong colocalization of GFP-TAZ (green) and mCherry-CycT1 (red), resulting in the appearance of yellow puncta (Fig. 6A and movie S2), indicative of coexistence within the same phase-separated compartments. Time-lapse analysis of the live cells expressing both GFP-TAZ and mCherry-CycT1 revealed that these condensates displayed phase separation behavior, including Brownian-like movement, shape remodeling, and fusion-driven growth. Furthermore, smaller puncta were frequently captured by larger, slower-moving condensates, leading to progressive enlargement and morphological maturation (Fig. 6B and movie S2). These findings suggest that TAZ has a dominant role in nucleating phase-separated condensates and can actively recruit CycT1 into shared nuclear compartments. Interestingly, co-immunoprecipitation assays showed that TAZ and CycT1 did not interact with each other directly (Fig. 6C), indicating that their co-localization within condensates is likely driven by multivalent interactions with shared nuclear components or through fusion of independently formed, nascent TAZ and CycT1 condensates. Indeed, as TAZ levels increased, the number of TAZ–CycT1 puncta decreased, while their average size increased (Fig. 6D). This shift is consistent with condensate growth through fusion of smaller condensates. Thus, TAZ promotes Cyc T1 phase separation in a dose-dependent manner and the co-localization of TAZ and CycT1 is mediated at least in part by fusion of the TAZ condensates with nascent CycT1 condensates.

**Fig. 6.**
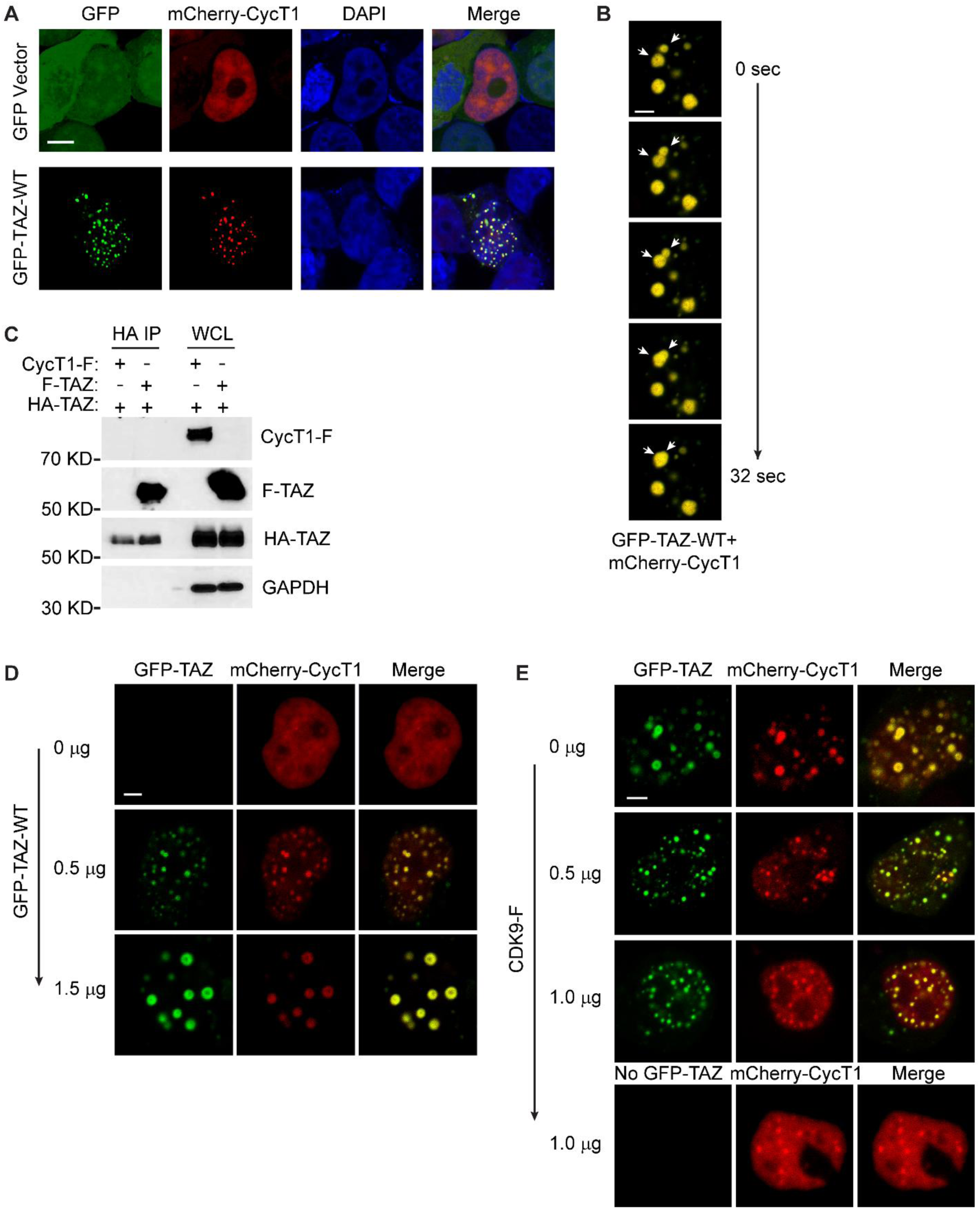
CycT1 and CDK9 effects on GFP-TAZ condensates. **(A)** Confocal images of HeLa cells 24 h after transfection by mCherry-CycT1 with empty vector or with GFP-TAZ-WT. Images were taken using Zeiss LSM 710 confocal microscope. The nuclei were stained by DAPI. **Bar**, 10 µm. **(B)** Live-cell confocal time series of a representative HeLa cell coexpressing GFP-TAZ-WT and mCherry-CycT1, acquired over 32 s. Arrows mark puncta undergoing fusion. **Bar**, 4 µm. **(C)** CycT1 does not directly bind to TAZ. 293T cells transfected with the indicated constructs were subjected to immunoprecipitation with anti-HA (HA IP), followed by western blotting with anti-Flag and anti-HA antibodies. GAPDH serves as a loading control. While CycT1-F did not precipitate with HA-TAZ, a positive control showed co-precipitation of Flag-TAZ (F-TAZ) with HA-TAZ. **(D)** TAZ dosage controls CycT1 partitioning. HeLa cells were transfected with mCherry-CycT1 together with increasing amounts of GFP-TAZ-WT (0, 0.5, 1.5 µg). **Bar**, 4 µm. **(E)** CDK9 modulates the organization of TAZ-CycT1 condensates. HeLa cells were co-transfected with fixed amounts of mCherry-CycT1 (1.0 µg) and GFP-TAZ (1.0 µg) together with increasing amounts of CDK9-F (0, 0.5, 1.0 µg). The bottom row (no GFP-TAZ) is a control lacking GFP-TAZ, with co-transfection of mCherry-CycT1 (1.0 µg) and CDK9-F (1.0 µg). **Bar**, 4 µm. The images in panels B, D and E were acquired on a Zeiss LSM880 Live-Cell confocal microscope. All confocal images are representative of at least three independent experiments.

Since CycT1 only exists in the complex with CDK9 as P-TEFb, and CDK9 has been shown to colocalize with CycT1 in phase-separated nuclear compartments (*16*), we next examined the impact of CDK9 on the formation and organization of TAZ–CycT1 condensates by live-cell confocal imaging. The experiments employed HeLa cells coexpressing GFP-TAZ and mCherry-CycT1, with or without ectopic expression of Flag-tagged CDK9 (CDK9-F). As shown in Fig. 6E, increased expression of CDK9 led to a pronounced reconfiguration of TAZ–CycT1 condensates, characterized by a higher number of nuclear puncta with reduced average size. This was accompanied by an increase in diffuse nuclear CycT1 signal, suggesting altered condensate dynamics. Notably, in the absence of GFP-TAZ, CDK9 overexpression alone was sufficient to promote the assembly of mCherry-CycT1 into punctate nuclear structures, although these appeared less distinct and more heterogeneous than condensates formed in the presence of TAZ. These observations indicate that CDK9 facilitates and stabilizes CycT1 phase separation and further modulates the spatial distribution and biophysical properties of TAZ–P-TEFb assemblies.

We next performed FRAP beam-size analysis to determine whether CycT1 alone or together with CDK9 affect the biophysical properties of TAZ condensates (Fig. 7A-C). First, we measured the effects of coexpressing mCherry-CycT1 on the dynamics of GFP-TAZ condensates (for images, see Fig. 6A). As shown in Fig. 7A, CycT1 induced a significant reduction in *R_f_* of GFP-TAZ measured with either objective (×40 or ×63). This suggests an increase in the fraction of immobile GFP-TAZ-WT molecules. Concomitantly, CycT1 expression reduced the τ values of GFP-TAZ by an order of magnitude (Fig. 7B). Of note, this led to a strong deviation of the τ(×40)/τ(×63) ratio from the 2.28 beam-size ratio expected for recovery by lateral diffusion (Fig. 7C), yielding a τ ratio of 1.3, close to the value of 1 which reflects recovery by exchange (*29, 30*). Thus, CycT1 expression results in a striking shift of the fluorescence recovery mode, from recovery by diffusion to recovery mainly by exchange, suggesting that under these conditions the GFP-TAZ-WT condensates display properties that resemble a shift towards a gel/solid-like state.

**Fig. 7.**
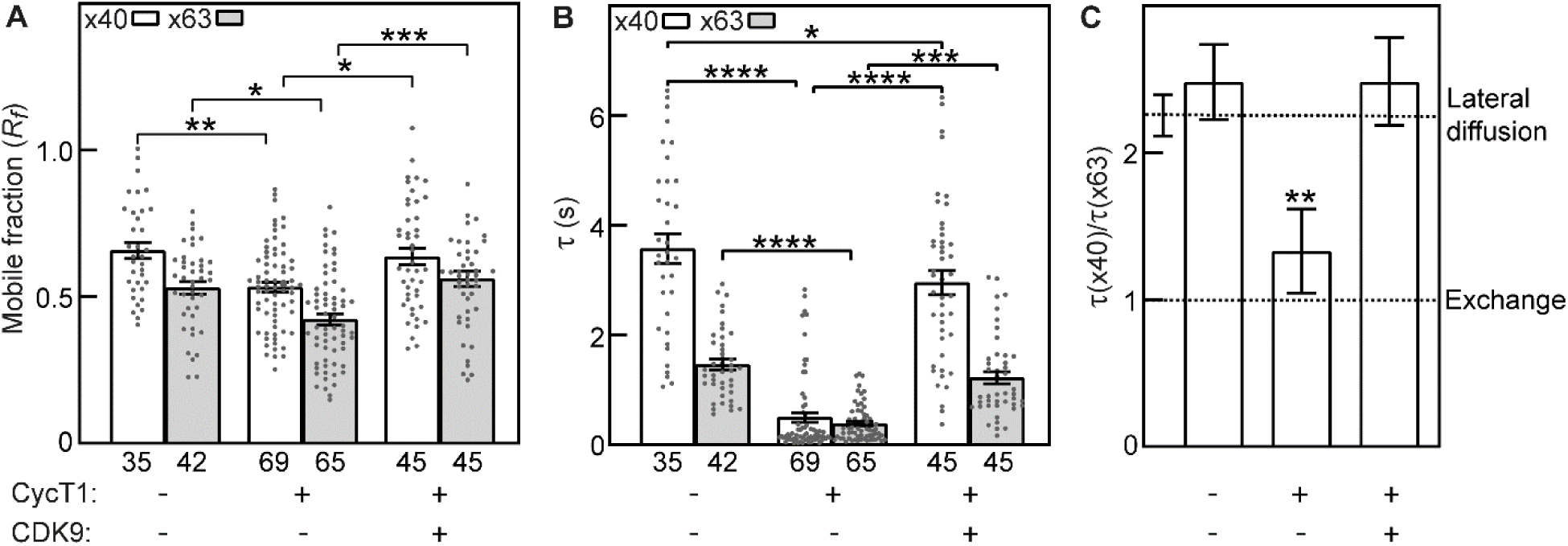
CycT1 shifts GFP-TAZ-WT condensates from recovery by diffusion to recovery mainly by exchange, and CDK9 reverses this effect. HeLa cells were transfected with GFP-TAZ-WT alone, or together with mCherry-CycT1 without or with CDK9-Flag. **(A, B)** FRAP beam-size analysis of the effects of CycT1 or CycT1 together with CDK9-Flag on GFP-TAZ-WT dynamics in nuclear condensates. Experiments were conducted 24 h post-transfection, as in Fig. 1. Bars are mean ± SEM of multiple independent measurements (the numbers of measurements are depicted under the bars). Asterisks represent significant differences between the indicated pairs (*, *P*<0.05; **, *P*<0.01; ***, *P*<10^−3^; ****, *P*<10^−4^; one-way ANOVA and Tukey’s post hoc test). **(C)** Bootstrap analysis of the τ(×40)/τ(×63) ratio relative to the beam size ratio (2.28). The SEM values for the *τ* ratios were calculated from the *τ* values shown in panel B, using 1,000 bootstrap resampling values. Asterisks indicate a significant deviation from the beam size ratio (**, *P*<0.01; Student’s two-tailed *t*-test).

Interestingly, the addition of CDK9 along with CycT1 led to a significant reversal of the effect of CycT1 alone on the FRAP of GFP-TAZ-WT, raising back the *R_f_* values, as well as the τ(×40) and τ(×63) values, close to those of singly-expressed GFP-TAZ-WT (Fig. 7A, B). Of note, the τ(×40)/τ(×63) ratio shifted back to the value expected for recovery by lateral diffusion (Fig. 7C). These results indicate that the inclusion of the complete transcription elongation factor P-TEFb shifts the properties of the GFP-TAZ biomolecular condensates to a fully liquid-like behavior. These results are in line with the imaging studies (Fig. 6), which showed a that the gel-like CycT1-TAZ condensates are larger than those of GFP-TAZ alone, while condensates obtained upon coexpression of GFP-TAZ with P-TEFb (CDK9 and CycT1) displayed a smaller average size.

### The ability of TAZ to form biomolecular condensates is closely associated with its biological functions

To determine the biological significance of TAZ phase separation, we examined two important aspects of the biological functions of TAZ using mammary epithelial cells and breast cancer as assay systems. TAZ is expressed at a high level in the mammary gland during postnatal development and plays essential roles in mammary epithelial cell differentiation and expansion during pregnancy (*39*), which is mimicked by a three-dimensional (3D) acinar morphogenesis model (*40*). In this model, untransformed MCF10A cells underwent proper morphological differentiation to form well-organized and polarized multicellular acini. Of note, knocking out (KO) TAZ by CRISPR-Cas9 significantly impaired acini formation, resulting in fewer cells per acinus and loss of apical-basal polarity in the 3D laminin-rich extracellular matrix (Fig. 8A, B). This defect was rescued by re-introduction of TAZ-WT, demonstrating the critical role of TAZ in acinar morphogenesis.

**Fig. 8.**
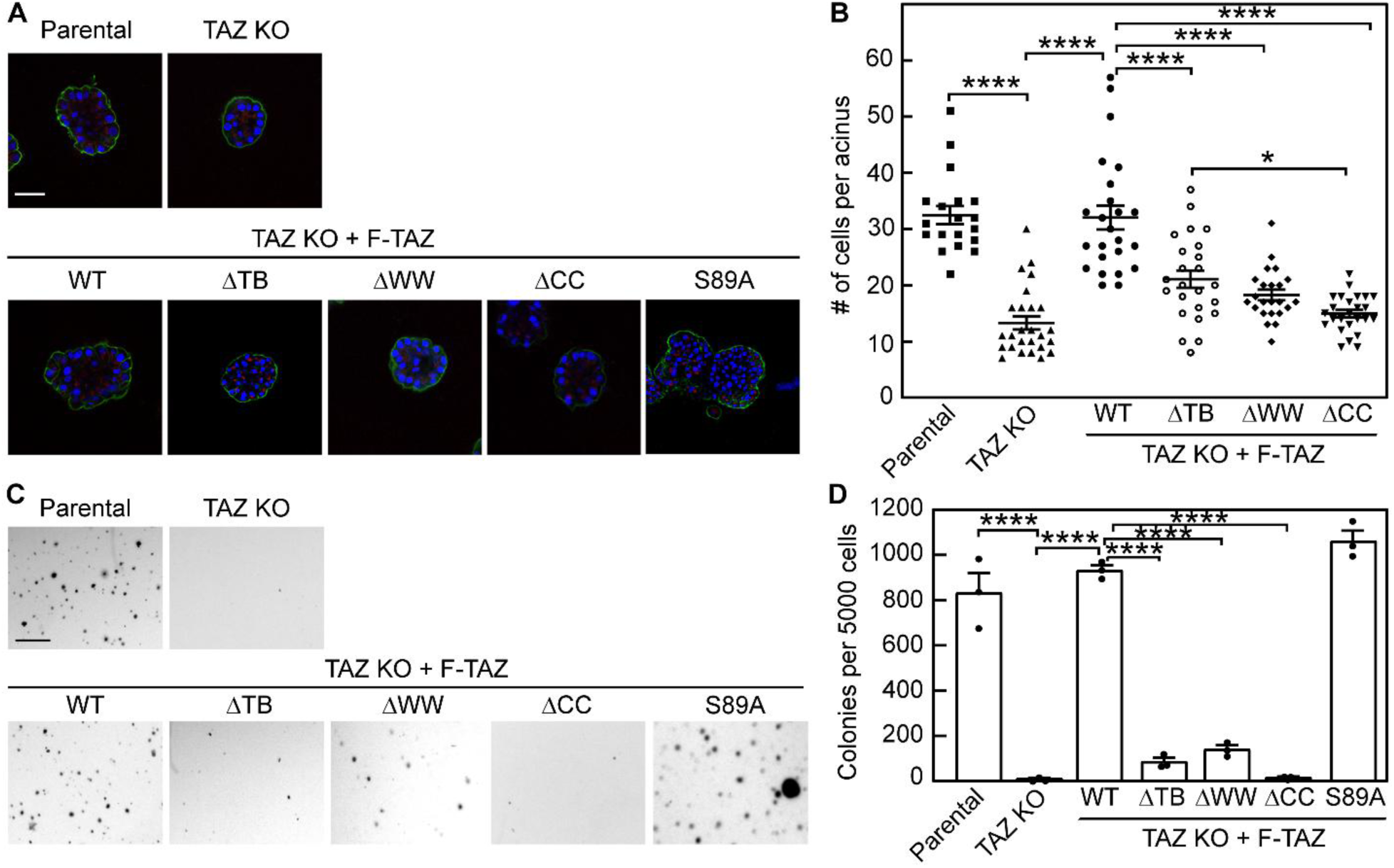
The ability of TAZ to undergo phase separation is closely associated with its biological functions. **(A)** 3D morphogenesis assays of MCF10A cells expressing Flag-TAZ variants. MCF10A TAZ KO cells were infected with the various Flag-tagged TAZ (F-TAZ) constructs as indicated. These cells, as well as the parental MCF10A cells, were cultured in 3D Matrigel for 7 days and stained for α-integrin (green) as a basolateral marker and GM130 (red) as an apical marker. Nuclei were counterstained with DAPI (blue). Representative confocal images were from three independent experiments. **Bar**, 40 μm. **(B)** Quantification of average acinar sizes based on the number of nuclei per acinus. A total of 20–25 acini from three independent experiments were analyzed per cell line. Asterisks depict significant differences between the pairs indicated by the brackets (*, *P*<0.05; ****, *P*<10^−4^; one-way ANOVA and Tukey’s post-hoc test). **(C)** Anchorage-independent growth in soft agar. MDA-MB-231 TAZ KO cells were infected with the designated F-TAZ constructs as indicated. Following infection, 5,000 cells were plated in 6-well soft agar plates and incubated for 21 days. Parental MDA-MB-231 cells and TAZ KO cells were used as controls. Colonies were stained with 1 mg/ml MTT. **Bar**, 100 μm. Colony numbers from each cell line expressing different TAZ constructs were counted and are shown in **(D).** Data are mean ± SEM of three independent experiments. Asterisks represent significant differences between the indicated pairs (****, *P*<10^−4^; one-way ANOVA and Dunnett’s post-hoc test).

TAZ is also a well-established oncogene frequently overexpressed in breast cancer, where its elevated levels are associated with increased malignancy and an enriched population of cancer stem cells (*41, 42*). Accordingly, MDA-MB-231 breast cancer cells, which express high levels of TAZ, exhibited strong anchorage-independent growth (Fig. 8C, D). In contrast, KO of TAZ from these cells significantly reduced colony formation, while re-introducing TAZ-WT into the TAZ KO cells restored their ability to form colonies (Fig. 8C, D), indicating that TAZ is essential for maintaining tumorigenic properties.

The ability of TAZ to form biomolecular condensates is closely associated with its biological function, as domain deletions that disrupt phase separation also impair its ability to support 3D differentiation or its oncogenic activity. As shown in Fig. 8A-D, deletion of the WW domain (ΔWW), which reduced phase separation, led to impaired acini growth and decreased anchorage-independent colony formation. Deletion of the coiled-coil domain (ΔCC), which completely abolished phase separation, resulted in a loss of both acini formation and anchorage-independent growth. In contrast, the constitutively active TAZ-S89A mutant, which exhibited enhanced phase separation (Figs. 2 and 3) and elevated transcriptional activation (*25*), formed disorganized and polarity-disrupted acini (Fig. 8A), while displaying robust anchorage-independent growth (Fig. 8C, D), consistent with its ability to induce oncogenic transformation (*42, 43*). Interestingly, the ΔTB mutant that formed only unstable puncta failed to support 3D acini growth and anchorage-independent colony formation, suggesting that more mature and stabilized TAZ puncta are critical for its function. Taken together, these results demonstrate that the ability of TAZ to undergo phase separation, regulated and stabilized by its structural domains, is a key determinant of its activities in cellular morphogenesis and tumorigenic potential.

## DISCUSSION

Phase separation has been recognized as an important mechanism that is essential for the activation and regulation of many biochemical reactions within the cells (*6–8*). While a growing number of cellular proteins have been shown to undergo phase separation, relatively little is known about the mechanism of assembly and maturation of these condensates, or the identities and varieties of condensates in the cells, mostly due to technical constraints. In this study, we investigated the dynamic assembly of TAZ biomolecular condensates using FRAP analysis with two laser beam sizes, along with quantitative fluorescence microscopy and cell biology methods. Our results showed that TAZ biomolecular condensates are multiphasic, more stable in the center and more labile at the periphery. These condensates initiate as small nascent clusters by self-nucleation through the TAZ CC domain based on disulfide bond formation involving C262, and gradually transition into larger more mature condensates. The TAZ CC domain is absolutely required for such transition, while the TAZ WW domain has a role in mediating multivalent protein interactions, which support the formation of the large visible condensates. This time-dependent maturation process is further stabilized by interaction with transcription factors and complexes including TEAD4 and P-TEFb. We further showed that the ability of TAZ to form large stable and mature biomolecular condensates is a key determinant of its activities in cellular morphogenesis and tumorigenesis. Our study presents a detailed mechanistic analysis of the intermediate steps of TAZ phase separation and reveals a highly dynamic nature of these biomolecular condensates in their maturation and functional activation.

TAZ functions as a scaffold that drives condensate formation and recruits client molecules such as TEAD4 and P-TEFb, which in turn influence the behavior or properties of the TAZ condensates (*25*). In line with earlier studies (*25, 44, 45*), we find that TAZ-WT forms visible condensates in the cell nucleus; these condensates display the behavior expected for liquid-liquid phase separation, as FRAP recovery occurs by diffusion in both phases (within the condensate at a slow rate, and in the surrounding nucleoplasm at a fast rate) (Fig. 1). These condensates displayed multiphasic properties, being more stable at the center (lower mobile fraction) and more labile at the periphery (Figs. 1, 2). Studies on the time-dependent maturation of TAZ variants (WT, S89A, ΔWW and ΔCC) showed early formation of TAZ nascent nucleoplasmic clusters, which can be detected by super-resolution imaging (Figs. 2, 3 and figs. S1, S2). The nascent clusters are especially prominent for TAZ mutants defective in the formation of visible condensates (ΔCC and ΔWW). TAZ-ΔCC cannot proceed beyond the nascent clusters stage, forming only nascent clusters that are far more labile (showing much faster exchange) than those formed by TAZ-WT or TAZ-S89A. This indicates an absolute requirement for the TAZ CC domain for the formation of proper initial clusters which allow multivalent interactions to recruit additional components for stabilization and growth into visible condensates. On the other hand, TAZ-ΔWW also forms a high level of labile initial clusters, which unlike the ΔCC mutant can mature to visible condensates (albeit smaller and more labile than those of TAZ-WT) at later times (72 h) (Figs. 2, 3; figs. S1, S2). We propose that the TAZ condensates have a multiphasic architecture, meaning that molecules in the center and in the periphery of the condensate are different in properties and possibly in composition. The center part may form mainly by self-assembly based on the TAZ CC domain and its C262 residue (Fig. 4) and on interactions with some additional specific components, while the peripheral region may contain additional molecules drawn to the condensate phase through interaction with the TAZ WW or TB domains. These multivalent interactions increase the concentration of molecules in the condensates and may stabilize the phase behavior, allowing the condensates to grow in size and gain functional capacities.

Of note, the addition of new components to the TAZ condensates can potentially alter their size and properties, rendering them capable of acquiring new gel/solid-like, or liquid-like features, as well as altering the stability and/or the functional state of the condensates. One example is the recruitment of TEAD into TAZ condensates, which leads to their stabilization, as shown by the reduced mobile fraction of TAZ in the presence of TEAD, and the lower stability of TAZ-ΔTB condensates either in the absence or presence of overexpressed TEAD (Fig. 5). A striking example of the flexibility of the condensates is given by the effects of fusion between TAZ condensates and P-TEFb condensates (Fig. 6 and movie S2). Here, expression of Cyc-T1 alone reduced *R_f_* of TAZ-WT and shifted its FRAP mechanism towards exchange, indicating slower diffusion within the condensates such that the exchange rate prevails, a behavior resembling a gel-like phase. However, coexpression with both Cyc-T1 and CDK9 reverted the dynamics back to those characteristics of liquid-liquid phase separation (Figs. 6, 7).

Given that the multivalent interactions within the condensates are dynamic and reversible, the condensates are likely to be in a constant state of assembly and disassembly, endowing them with the flexibility required to meet different functional demands in response to upstream signals. Thus, in a given cell, TAZ biomolecular condensates are likely to be highly heterologous with different compositions and in different functional and regulatory states.

The current studies employed expression of transfected tagged TAZ proteins in cells to provide proof of concept for the mechanisms of condensate formation and maturation. However, since we are examining the internal structure of the condensate, the impact of overexpression is less of a concern, especially as the TAZ condensates appear to be heterogenous with respect to interacting partners and biological functions. In addition, some of the condensates may be subject to regulation by upstream ligands. It would be interesting to examine the effects of specific proteins that interact with the TAZ WW domain on condensate formation, co-localization and maturation. These proteins could potentially serve as markers for specific TAZ condensates.

One unsolved question in the field of phase separation is how condensates containing multiple proteins or complexes are formed; are additional proteins recruited to the condensates through direct protein-protein interaction or through fusion with other condensates that contain additional client molecules? Our results showed that both mechanisms can be involved. In the case of TEAD4, its recruitment to the TAZ condensates depends on the physical interaction between TEAD4 and TAZ, because TAZ mutants defective in TEAD4 binding failed to recruit TEAD4 into the condensates. In contrast, P-TEFb does not directly interact with TAZ but forms its own nuclear condensates. Its co-localization with the TAZ condensates is achieved through fusion of the P-TEFb and TAZ condensates. This fusion process is dynamic and reversible, with smaller puncta fused into bigger puncta and big puncta disappearing later. The recruitment of these additional molecules into the TAZ condensates further stabilizes the condensates and enables their transcriptional activity (Fig. 6 and Ref. (*25*)). This ability of TAZ to phase separate is critical for efficient and TAZ-specific gene expression (*25*). In support of this model, we found that mutations in TAZ disrupting phase separation resulted in defective morphological differentiation of mammary epithelial cells in a 3D acinar differentiation model and abolished its oncogenic activity in breast tumorigenesis (Fig. 8). As TAZ, but not the closely related YAP, plays an essential role in mammary gland development and breast cancer progression (*39, 46*), this ability to phase separate provides an effective mechanism to insulate TAZ signaling from that of YAP, allowing TAZ-specific transcription and biological activities. Thus, phase separation could be a mechanism that is generally utilized by all signaling pathways and molecules to generate signaling and regulatory specificity.

## MATERIALS AND METHODS

### Reagents

Dulbecco’s modified Eagle’s medium (DMEM; cat. #01-052-1A) and cell culture reagents (fetal calf serum-FCS, L-glutamine, penicillin-streptomycin) were from Biological Industries Israel (Beit Haemek, Israel-Sartorius group) or Gibco/Thermo Fisher Scientific (Waltham, MA). DMEM/F12 medium (cat. #11320-033), Opti-MEM (cat. #11058021) and horse serum (cat. #16050-122) were from Gibco. Hanks’ balanced salt solution (HBSS) with Ca^2+^/Mg^2+^ without phenol red (cat. #009015237500) and 4-(2-hydroxyethyl)-1-piperazineethanesulfonic acid (HEPES, 1 M, pH 7.3; cat. #000773233100) were from Bio-Lab Ltd. (Jerusalem, Israel). Opti-MEM (cat. #11058021) was from Gibco. Lipofectamine^TM^ 3000 (cat. #L3000001) was from Invitrogen-Thermo Fisher Scientific (Waltham, MA). Insulin (cat. #1-1882), cholera toxin (cat. #C8052) and hydrocortisone (cat. #H-0888) were from Sigma-Aldrich (St. Louis, MO). Human EGF (cat. #AF-100-15-500UG) was from PeproTech (Rocky Hill, NJ).

### Antibodies

Anti-α6-integrin (rat monoclonal NKI-GoH3, cat. #MAB1378) was from Chemicon/Sigma-Aldrich. Murine monoclonal anti-Golgi matrix protein of 130 (GM130; cat. #610823) was purchased from BD Pharmingen (San Diego, CA), and murine anti-a-tubulin (cat. #ab7291) from Calbiochem (San Diego, CA). Murine monoclonal anti-Flag (M2; cat. #F1804) and anti-HA (12CA5; ROAHA) antibodies were from Sigma-Aldrich. Horseradish peroxidase (HRP)-conjugated goat anti-rabbit (cat. #111-035-144) and anti-mouse (cat. #115-035-003) antibodies were from Jackson ImmunoResearch Laboratories (West Grove, PA). Goat anti-rat IgG (H+L) labeled with Alexa Fluor 488 (cat. #A-11006) and Goat anti-mouse IgG (H+L) labeled with Alexa Fluor 546 (cat. #A-11030) were from Invitrogen/Thermo Fisher Scientific.

### Plasmids

Expression vectors of GFP-TAZ and of GFP-TAZ mutants with deletion of the WW domain (GFP-TAZ-ΔWW) or the CC domain (GFP-TAZ-ΔCC) in the pEGFP-C1 vector (TakaraBio-Clontech, Mountain View, CA), as well as of the constitutively active GFP-TAZ-S89A mutant, were described earlier (*25*). The GFP-TAZ mutant lacking the TEAD binding domain (GFP-TAZ-ΔTB) was generated by direct PCR using the KAPA HiFi PCR Kit (Roche) with the following primers: forward, 5′-CTGTACAAGTCCGGACTCAGATCTCGAGCTCTGCCCCCGGGCTGGGAGA-TGACCTTCACG-3′; reverse, 5′-CGTGAAGGTCATCTCCCAGCCCGGGGGCAGAGCTCG-AGATCTGAGTCCGGACTTGTACAG-3′. The full-length pEGFP-C1-TAZ plasmid was used as template. The expression vector pmCherry-CycT1 was constructed by replacing the EGFP coding sequence in pEGFP-C1 with mCherry. The vector pCMV5B was used for expression of Flag-TEAD4, CDK9-FlagFlag-TAZ and HA-TAZ. The vector pRK5 was used for expression of CycT1-Flag.

TAZ-CC domain (WT, or containing the C262S mutation) fused to the TRX-His_6_-tag were generated *via* standard PCR-based methods and inserted into pET-based vectors containing TRX-His_6_-tag followed by an HRV-3C protease cleavage site. The recombinant protein sequence information was as follows: WT TAZ-CC, 201P-270E, Q9GZV5 (Uniprot); the Ser mutation of TAZ-CC Cys262 was introduced by site-directed mutagenesis using the following primers: forward, 5’-AAGCTGCCCTCAGTCGACAGCTCCCCATGGAAG-3’; reverse, 5’-TCGACTGAGGGCAGCTTCCTGCCTCATGAGCT-3’.

### Protein expression and purification

Plasmids containing the TRX-His_6_-tag gene were transformed into Escherichia coli Rosetta (DE3) cells using the heat-shock method. Cells were grown in LB medium at 37℃ to an optical density at 600 nm of 0.6-0.8. The expression of recombinant proteins was induced by 0.2 mM isopropyl-β-D-thiogalactoside (IPTG; cat. #MB3026, MeilunBio) and 0.01 mM ZnCl_2_ at 16℃ for 18 h. Each recombinant protein was purified using a nickel-NTA agarose affinity column (cat. #30210, Qiagen, Hilden, Germany) followed by size-exclusion chromatography (Superdex 75, cat. #29148721, Cytiva, Marlborough, MA) with a column buffer containing 50 mM HEPES, pH 7.5, and 100 mM NaCl.

### Cell culture and transfections

HeLa cells (cat. #CCL-2), MCF10A human mammary epithelial cells (cat. #CRL-10317), MDA-MB-231 human breast cancer epithelial cells (cat. #HTB-26) and 293T human kidney epithelial cells (cat. #CRL-3216) were from American Type Culture Collection (ATCC, Manassas, VA). Hela cells and 293T cells were grown in DMEM supplemented with 10% FCS, 1% penicillin/streptomycin and 2 mM L-glutamine. MCF10A cells were cultured in DMEM-F12 supplemented with 5% horse serum, 20 ng/ml EGF, 10 µg/ml insulin, 0.5 µg/ml hydrocortisone, 100 ng/ml cholera toxin, and 1% penicillin/streptomycin. MDA-MB-231 cells were cultured in DMEM supplemented with 10% FCS and 1% penicillin/streptomycin. All cell lines were grown at 37°C with 5% CO_2_. They were authenticated at UC Berkeley Cell Culture Facility by single nucleotide polymorphism testing. The HeLa cell line was also authenticated by STR analysis at the Genomics Center of the Biomedical Core Facility, Technion, Haifa, Israel. The cells were routinely analyzed by RT-PCR for mycoplasma contamination and found to be clean.

For FRAP or fluorescence microscopy studies, HeLa cells grown on glass coverslips in 6-well plates were transfected by a total of 2.5 μg DNA of expression vectors for GFP-TAZ (WT or specific mutants; 1.3 μg) alone or together with expression vectors for TEAD4 or CycT1 (1.2 μg) using Lipofectamine^TM^ 3000 according to the manufacturer’s instructions. Cells were subjected to studies by fluorescence microscopy or FRAP beam-size analysis (described below) at 12-72 h post-transfection, as mentioned in the figure legends.

For protein-protein interaction assays, 5.0 μg of pCMV5B-HA-TAZ, together with 5.0 μg of pRK5-CycT1-Flag or pCMV5B-Flag-TAZ, were transfected into 3.5×10^6^ 293T cells using Lipofectamine^TM^ 3000. 48 h post-transfection, cells were harvested and subjected to immunoprecipitation followed by Western blotting as described (*47*).

### Size-exclusion chromatography coupled with multi-angle light scattering assay

The size exclusion chromatography coupled with multi-angle light scattering (SEC-MALS) system is composed of a static light scattering detector (miniDawn, Wyatt Technology, Santa Barbara, CA), a differential refractive index detector (Optilab, Wyatt Technology), and an AKTA purifier (GE Healthcare, Chicago, IL). 100 μL sample was injected into a Superdex 200 Increase 10/300 GL column (cat. #28990944, Cytiva) pre-equilibrated with a column buffer containing 50 mM HEPES, pH 7.5, and 100 mM NaCl. Data were analyzed by the ASTRA6 (Wyatt Technology) software and the fitted molecular weight was obtained.

### FRAP and FRAP beam-size analysis studies

HeLa cells transfected with a *GFP-TAZ* variant were subjected to FRAP and FRAP beam-size analysis experiments as described by us earlier (*25, 29–31*). Experiments were conducted at 22 °C in HBSS supplemented with 20 mM HEPES, pH 7.2. For measurements using a smaller Gaussian laser beam size, an argon-ion laser beam (Innova 70C, Coherent, Santa Clara, CA) was focused *via* an AxioImager.D1 microscope (Carl Zeiss MicroImaging, Jena, Germany) equipped with a plan-apochromat oil immersion objective (×63/1.4 NA) to a Gaussian spot with a radius *ω* = 0.77 ± 0.03 μm. For measurements with a larger Gaussian beam, a C apochromat water-immersion objective (×40/1.2 NA) was used, resulting in a Gaussian radius of *ω* = 1.17 ± 0.05 µm. In this setup, the ratio between the illuminated areas, *ω*^2^(×40)/*ω*^2^(×63), was 2.28 ± 0.15 (*n* = 59; SEM of the ratios calculated from the bootstrap values as described below). The fluorescence intensity at the beam-illuminated spot was measured with the laser beam at monitoring intensity (488 nm, 1 μW), followed by a brief 5 mW pulse (7–14 ms) that bleached 60–75% of the fluorescence in the illuminated spot. Fluorescence recovery was followed using the monitoring beam intensity. The apparent characteristic fluorescence recovery time (*τ*) and mobile fraction (*R_f_*) for each curve were derived by nonlinear regression analysis, fitting to a lateral diffusion process (*29*). The significance of differences between sets of τ or *R_f_* values measured with the same beam size was evaluated by one-way ANOVA and Tukey’s post hoc test. To compare the *τ*(×40)/*τ*(×63) ratio of a TAZ-GFP mutant with the *ω*^2^(×40)/*ω*^2^(×63) beam-size ratio, statistical significance was evaluated by unpaired two-tailed Student’s *t*-test using bootstrap analysis with 1,000 bootstrap samples (*48*), which is preferable to compare between ratios, as described by us earlier (*25, 49*).

### Fluorescence imaging and super-resolution microscopy

Cells grown on glass coverslips were transfected with expression vectors for *GFP-TAZ* variants as described under “Cell culture and transfections”. At various time points post transfection, they were fixed with 4% paraformaldehyde in PBS (20 min, 22 °C), washed by PBS and mounted with DAPI-containing mounting medium (Cat. #AB104139, Abcam, Cambridge, UK) and sealed. For confocal imaging, fluorescence was detected using a Zeiss LSM 710 confocal microscope or Zeiss Elyra PS1 super-resolution structured illumination microscope. For live cell imaging, images were taken by Zeiss LSM880 Live-Cell confocal microscope and the videos made from collected images were processed by Imaris Software (Oxford Instruments, Abingdon, Oxfordshire, England).

To measure the time-dependent formation of nascent GFP-TAZ nucleation centers at the initial stages of forming visible condensates, we conducted super-resolution microscopy experiments on HeLa cells transfected and prepared as above, and measured the density of nascent puncta (below the resolution of regular light microscopy) in their nuclei, as well as the nuclear density of puncta visible at standard fluorescence microscopy resolution (visible condensates). To this end, we employed a Zeiss LSM800 Airyscan super-resolution microscope with a ×63/1.4 NA oil immersion objective, followed by Airyscan processing (2D, default settings). Puncta in the nuclei of multiple cells were counted by an in-house Fiji (Image J) script, with a threshold for nascent puncta set at 0.008 μm^2^ (diameter of 0.1 μm, the resolution of the Airyscan microscope) and an upper limit of 0.05 μm^2^ (diameter of ∼0.24 μm, above which the puncta are visible in standard fluorescence microscopy). To measure the nuclear density of visible puncta (*i.e*., condensates), a lower threshold of 0.06 μm^2^ (diameter of 0.28 μm) was set. The counting was performed on the nuclear regions, and normalized per the area size of each nucleus.

### Soft agar assay

Anchorage-independent growth assay was performed as described previously (*39*). 5,000 cells were suspended in 2 ml DMEM/10% FCS) containing 0.44% Bacto Agar (Becton Dickinson) and overlaid on the hardened 0.66% agar bottom layer in a well of a 6-well cluster. Fresh medium (1 ml) was added to each well once a week for 3 weeks. The colonies were visualized by staining with 0.5 mg/ml 3-(4,5-dimethylthiazol-2-yl)-2,5-diphenyl tetrazolium bromide (MTT; cat. #M5655, Sigma-Aldrich) for 4 h at 37°C.

### Three-Dimensional Morphogenesis Assay

Differentiation of MCF10A cells in 3D laminin-rich extracellular matrix (lrECM) was performed as described (*39, 40*). Cultrex basement membrane extract (BME) (cat. #3433-005-02, Trevigen, Gaithersburg, MD) was used as a substitute for Matrigel. Microscopy was performed on a Zeiss LSM710 confocal microscope at the UC-Berkeley Biological Imaging Facility. The images were generated by serial confocal cross sections.

### Statistical analysis

Statistical analysis was done by Prism10 (GraphPad Software, San Diego, CA). Significant differences between multiple data sets were evaluated by one-way ANOVA followed by the relevant post-hoc test (Tukey’s or Dunnett’s). Student’s *t*-test was used to calculate the significance of the difference between two groups. Data are presented throughout as mean ± SEM, along with the number of independent measurements (given within each figure or figure legend). FRAP beam-size ratio measurements used bootstrap analysis with 1000 bootstrap samples (*48*), as described previously (*25, 49*). *P* values below 0.05 were defined as statistically significant. All attempts at replication were successful with similar results.

## Acknowledgments

We thank Denise Schichnes and Juliana Cho at the CNR biological imaging facility at the University of California, Berkeley for assistance with microscopy.

## Funding

This research was supported by grant no. 2110314 from the U.S. National Science Foundation (NSF) and by grant no. 2020700 from the U.S.-Israel Binational Science Foundation (BSF) to KL and YIH.

## Author contributions

Conceptualization: YIH, KL, KES

Methodology: KES, QZ, EG, HW, XDL

Investigation: KES, QZ, HW, EG, XC, YJ

Visualization: KES, QZ, YIH, KL, XC, YJ, XDL

Supervision: YIH, KL

Funding acquisition: KL, YIH

Writing - original draft: YIH, KL, KES

Writing - review & editing: KL, YIH, KES, QZ, HW, EG, XDL, XC, YJ

## Competing Interests

Authors declare that they have no competing interests.

## Data and materials availability

All data generated or analyzed during this study are available in the main text or the supplementary materials.

## Notes

### Competing Interest Statement

The authors have declared no competing interest.

