## Supplementary materials for "Analysis of the assembly, stabilization and maturation of the multiphasic TAZ biomolecular condensates"

Keren E. Shapira *et al.*

**This PDF file includes:**

Figs. S1 to S2
Legends to Movies S1 to S2

**Other Supplementary Materials for this manuscript include the following:**

Movies S1 to S2

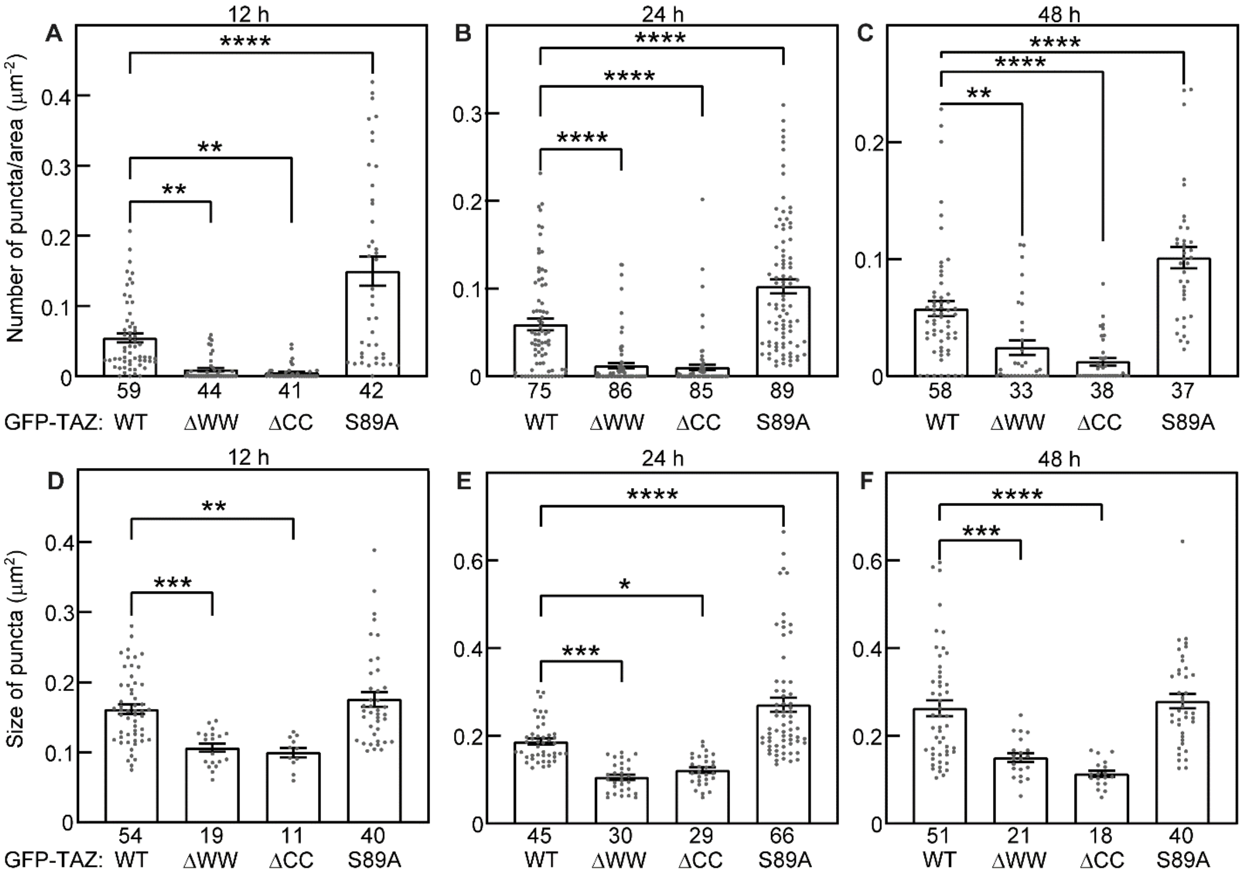


**Fig. S1.**

**Original data and statistics of the time course for visible condensates formation by GFP-TAZ-WT and mutants**. Super-resolution images of HeLa cells transfected with GFP-TAZ-WT, -S89A, -ΔWW or -ΔCC were analyzed as described in Fig. 3. **(A-C)** The number of visible puncta per area in the nucleus. (**D-F**) The size of visible puncta. Data are mean ± SEM. The time post-transfection is depicted above each panel. To measure only visible condensates, a lower threshold of 0.06 $\mu m^2$ was set for the counting by Fiji. The numbers of measurements (each on a different cell) are depicted underneath each bar. Significant differences between the indicated pairs were evaluated by one-way ANOVA and Tukey's post hoc test (*, $P$<0.05; **, $P$<0.01; ***, $P<10^{-3}$; ****, $P<10^{-4}$).

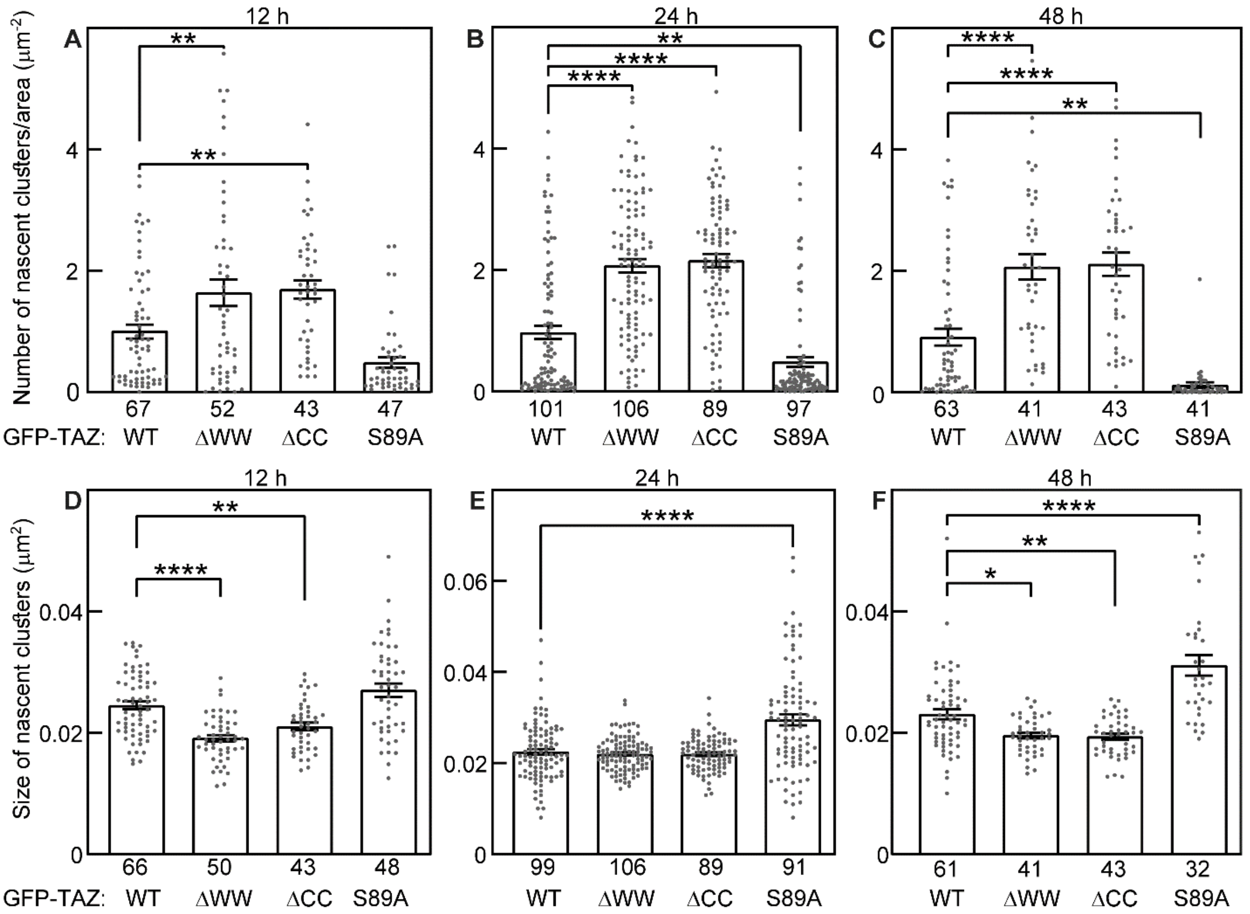


**Fig. S2.**

**Original data and statistics of the time course for nascent clusters formation by GFP-TAZ variants**. HeLa cells were transfected with GFP-TAZ-WT, -S89A, -DWW or -DCC and subjected to the super-resolution studies described in Fig. 3. **(A-C)** The number of nascent clusters per area in the nucleus. **(D-F)** The size of nascent clusters. Data are mean ± SEM. The time posttransfection is depicted above each panel. Nascent clusters are defined as clusters occupying an area between 0.008 $\mu m^2$ and 0.05 $\mu m^2$, counted by Fiji. The numbers of measurements (each on a different cell) are depicted underneath each bar. Significant differences between the indicated pairs were evaluated by one-way ANOVA and Tukey's post hoc test (*, $P$<0.05; **, $P$<0.01; ***, $P<10^{-3}$; ****, $P<10^{-4}$).

**Movie S1.**

**mCherry-CycT1 in live HeLa cells.** Live confocal recording of HeLa cells expressing mCherry-CycT1 alone. Images were taken by Zeiss LSM880 Live-Cell confocal microscope for 300 cycles over 5 min, to capture nuclear dynamics. The videos made from collected images were processed by Imaris Software. **Bar**, 2 μm.

**Movie S2.**

**GFP-TAZ recruits mCherry-CycT1 into nuclear condensates.** Live confocal recording of HeLa cells coexpressing GFP-TAZ and mCherry-CycT1. Images were acquired for 300 cycles over 5 min. The videos made from collected images were processed by Imaris Software. **Bar**, 2 μm.
